# Machine Learning-Assisted Pathway Optimization in Large Combinatorial Design Spaces: a p-Coumaric Acid Case Study

**DOI:** 10.1101/2025.06.13.659482

**Authors:** Paul van Lent, Rianne van der Hoek, Sara Moreno Paz, Irsan Kooi, Moniek Jonkers, Priscilla Zwartjens, Joep Schmitz, Thomas Abeel

**Affiliations:** Intelligent Systems, Delft University of Technology, van Mourik Broekmanweg, Delft, 2628XE, Zuid-Holland, Netherlands; Department of Science and Research, dsm-firmenich, Alexander Fleminglaan 1, City, 2613AX, Zuid-Holland, Netherlands; Infectious Disease and Microbiome Program, Broad Institute of MIT and Harvard, Main St., Cambridge, 02142, MA, United States of America

**Keywords:** Metabolic Engineering, Combinatorial Pathway Optimization, Strain Recommendation, Machine Learning

## Abstract

Combinatorial pathway optimization is an important tool for industrial metabolic engineering to improve titer, yield, or productivity of strains. Machine learning has been increasingly applied on many aspects of the Design-Build-Test-Learn (DBTL) cycle, an engineering framework that aims to navigate through the large landscape of theoretically possible designs using an iterative approach. While machine learning-assisted recommendation strategies have been successfully used to optimize strains, they have so far been limited to relatively small design spaces with few targeted elements. This small design space may limit key strengths of these approaches, such as strong predictive capabilities of supervised machine learning and exploration-exploitation schemes widely used in reinforcement learning and Bayesian optimization. In this work, two DBTL cycles are performed on *Saccharomyces cerevisiae* for p-coumaric acid production. We first perform a large library transformation on eighteen genes with twenty promoters, which expands the size of the combinatorial design space significantly (approximately 170 million configurations), followed by a smaller model-guided recommendation round. We use a machine learning-assisted recommendation strategy, based on the gradient bandit algorithm, parametrized to balance exploration and exploitation. We show that our recommendation strategy has a better performance than strain recommendation strategy using greedy strategies, such as feature importance-based methods. While balancing between exploration and exploitation has been shown to be important in many applications, we provide the first direct experimental illustration of this effect by recommending strains for scenarios with increasing exploitative-ness. A clear effect of the exploration-exploitation scenario on the p-coumaric acid production distribution of strains is observed, where a balanced scenario shows a higher variation in production over an exploratory or exploitative scenario. Interestingly, using an alternative top-producing parent strain with this balanced exploration-exploitation scheme gives the highest p-coumaric acid production, suggesting that model predictions outside of the training data distribution can still be used to perform successful strain recommendation. Overall, these results suggest that using machine learning-assisted strategies with balanced exploration-exploitation can be used to efficiently explore large combinatorial design spaces. The best engineered strain shows an increase in p-coumaric acid production of 137% over the parent strains and a 0.07g/g yield on glucose.

## 1 Introduction

Transforming the production of chemical compounds into sustainable alternatives is an important step to mitigate the climate and biodiversity crisis [1]. Microbial cell factories provide a sustainable alternative for the production of many useful compounds such as pharmaceuticals, feed additives, and bulk chemicals [2]. However, the long and expensive development of microbial cell factories from proof-of-principle strains hinders the widespread use for sustainable production [3, 4].

Advances in genome engineering and high-throughput screening have paved the way for combinatorial pathway optimization [5]. In combinatorial pathway optimization, multiple pathway elements are targeted simultaneously, with the goal of finding the optimum pathway configuration that maximizes product titer, rate, yield (TRY) values [6]. This approach addresses the limitation of missing global optima in sequential optimization [5], but may result in combinatorial explosion of the pathway configuration space. This consequently results in only sparsely sampling the set of all possible pathway configurations, regardless of how high-throughput the screening procedure is [7]. To mitigate this challenge, metabolic engineers often choose to limit the number of pathway elements simultaneously optimized [8] by using the Design-Build-Test-Learn (DBTL) cycle to navigate through the vast combinatorial landscape [9] (Fig. 1 A). In the DBTL cycle, an initial set of pathway designs is chosen (Design phase), introduced in a base strain (parent) and screened for production TRY values (Build and Test phase). Then, the relationship between pathway design and production is learned to inform the next Design phase (Learn phase). From here, the next set of designs are chosen and introduced in the same base strain or a new parent strain with higher baseline production. This approach is computationally reminiscent of a Markov decision process (Fig. 1B), where the strain can be viewed as a state, a pathway design as an action, and the production value determined by high-throughput screening as a reward (1B). The overall goal is to maximize the reward (TRY values) in the long term and finding strategies that achieve this goal is reminiscent of the reinforcement learning problem [10, 11].

**Fig. 1:**
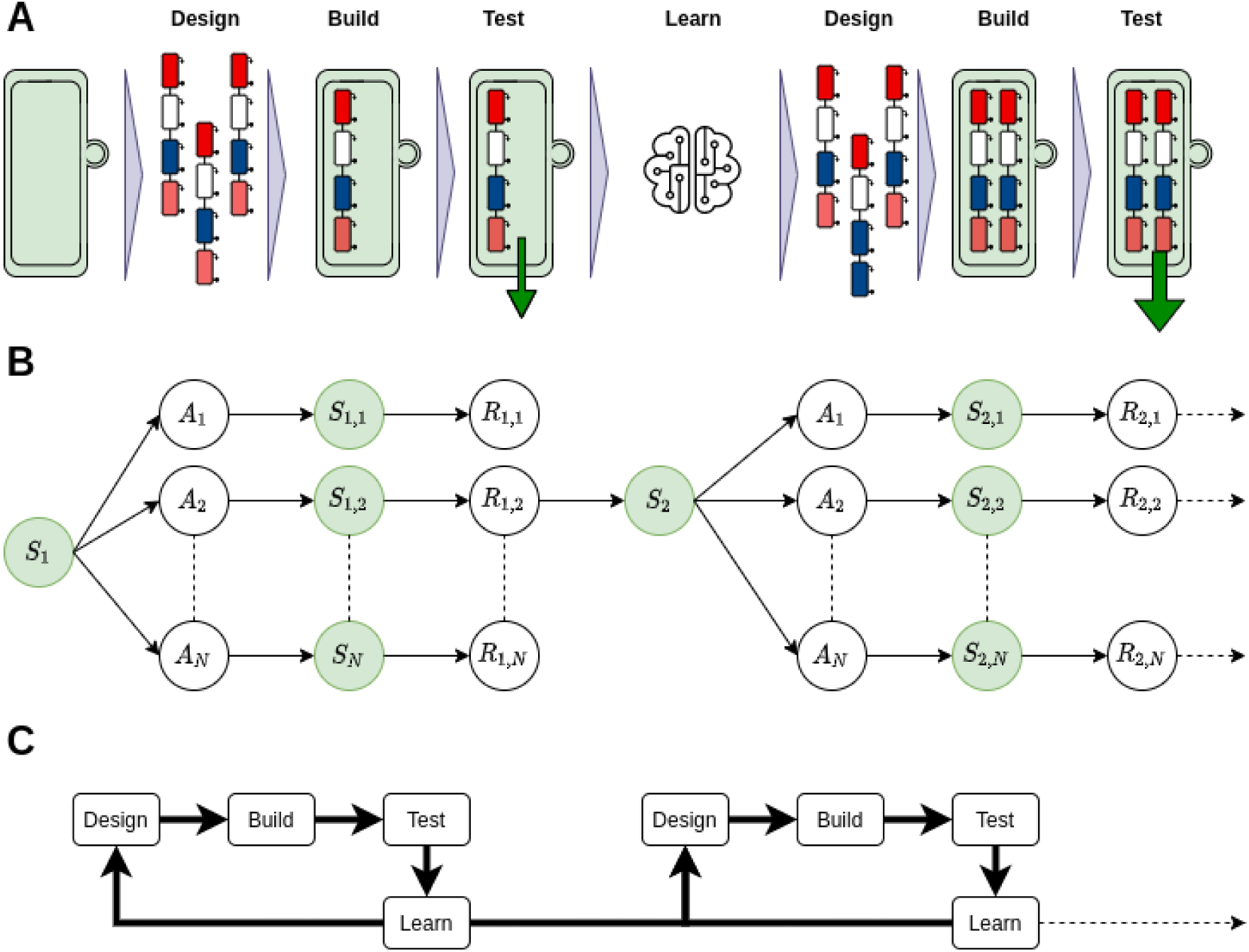
Iterative metabolic engineering from an experimental, computational, and metabolic engineering perspective. A) The parent strain is transformed using a set of pathway designs, which then integrates the pathway in its genome. This leads to a production of a compound of interest (green arrow). Using a set of built strains, a regressor can be trained to predict the pro- duction outcomes of new designs. This iterated until strains produce industrially relevant quantities, or until a theoretical maximum has been reached. B) Iterative metabolic engineering can be posed as a Markov Decision Process. Here, a subset of all actions *A* (designs) can be used to move to the next state *S* (new strains) that result in a reward *R* (production value). One can then either use supervised ML to learn what action was best given *S*_1_ or choose to move further in the MDP using that same model. The role of supervised ML-assisted automated recommendation is to optimally navigate through this Markov Decision Process. C) The DBTL cycle of metabolic engineering refers to the processes (arrows) when moving through the Markov Decision Process.

Machine learning has been increasingly used in the DBTL cycle, ranging from predicting design action outcomes [9], assessment of gene importance [12], and automated recommendation of new strains [13, 14]. Specifically, the recommendation of new designs is an exciting application with the potential to close the loop from Learn-to-Design and has shown some successful applications [15, 16]. However, there remains a limitation in how these machine learning recommendation strategies are applied. First, while advances in genome engineering and high-throughput screening have allowed for targeting many pathway elements simultaneously, the design configuration space often remains limited to a few genes. With configuration spaces in the order of thousands designs, high-throughput screening and sequencing can achieve a relatively dense sampling. This dense sampling results in potentially increasing production for the recommended strains only marginally, as strains that were built are already close to the optimum [17]. Another consequence of limiting the size of the combinatorial design space is that balancing the exploitation of the model prediction and exploration of new regions, a concept that is widely studied in Bayesian optimization and reinforcement learning, may not be important.

In this work, we investigate the influence of expanding the pathway configuration space significantly when recommending new designs. As a target, we choose p-coumaric acid (p-CA), which is an aromatic amino-acid-derived molecule from phenylalanine and tyrosine. p-CA is a precursor to many bio-active molecules with commercial applicability [18]. An initial large library that targets eighteen genes with twenty promoters resulting in approximately 170 million pathway design configurations, was transformed in *S. cerevisiae*. Despite the high-throughput capabilities, the set of designs sampled from these configurations is highly sparse (5 *×* 10*^−^*^6^%). We introduce a ML-assisted recommendation strategy, based on the gradient bandit algorithm [10], with XGBoost as a supervised model for predicting action outcomes [19]. Due to the high sparsity of the space, we experimentally investigate the effect of exploration-exploitation in this setting to find optimal designs. We further studied whether the supervised model can be used to predict action outcomes that are out of the training data distribution (i.e. when different parent strains are used), a scenario that is commonly encountered during iterative optimization. Our results suggest that a large library transformation round can be used to train a highly effective predictive model, which can be applied for recommending new strains. The balancing of exploration and exploitation is important for the trade-off between gene diversity of the recommended strains and p-CA production and is the preferred approach over greedy building of strains solely based on feature importance [12]. However, a combined approach that includes the generation of a new parental strain using a greedy approach, and a balanced exploration/exploitation that accounts for the increasing model uncertainty due to predicting actions outside the training data distribution, results in the highest performing strains. The best strain had an 137% improvement in p-CA production compared to the parent strain and a 0.07g/g yield on glucose.

## 2 Results

### 2.1 Library design using the *yeast8* genome-scale model

Supervised machine learning has been used for predicting TRY values of strains [12, 15, 20], but has typically been limited to small sets of genes and promoters. This hampers the strength of the strong predictive capabilities that machine learning offers, as strain performance may only marginally improve compared to the training data. We therefore propose to choose a larger set of gene targets and promoter values to significantly expand the combinatorial design space. To choose gene targets from thousands of reactions that are contained within the yeast metabolism, we used parsimonious flux balance analysis (pFBA) on the *yeast8* consensus genome-scale model [21, 22]. pFBA aims to find the minimal sum of fluxes that maximize the objective function.

The ratio of reaction fluxes between two pFBA objectives, p-CA and biomass, was used to select genes from reactions that divert flux from biomass to p-CA (Fig. 2A). This led to the identification of 33 targets after excluding sink, exchange and non-targetable reactions from the pFBA solution. Targets were validated for their effect on p-CA or other aspects of the aromatic amino-acid biosynthesis route, as well as thermodynamic considerations of the reactions (see B.1). This resulted in 18 genes to be considered in the DNA library, including two glycolysis genes (FBA, ENO2), seven pentose phosphate pathway genes (ZWF1, SOL3/4, GND1/2, RKI, TKL), eight elements of the aromatic amino-acid biosynthesis pathway (ARO1, ARO4, ARO2, PHA2, ARO8, PAL1, CPR1, C4H), and one gene involved in transfer of oxoglutarate and citrate between the mitochondrion and cytosol (YHM2) (Fig. 2B).

**Fig. 2:**
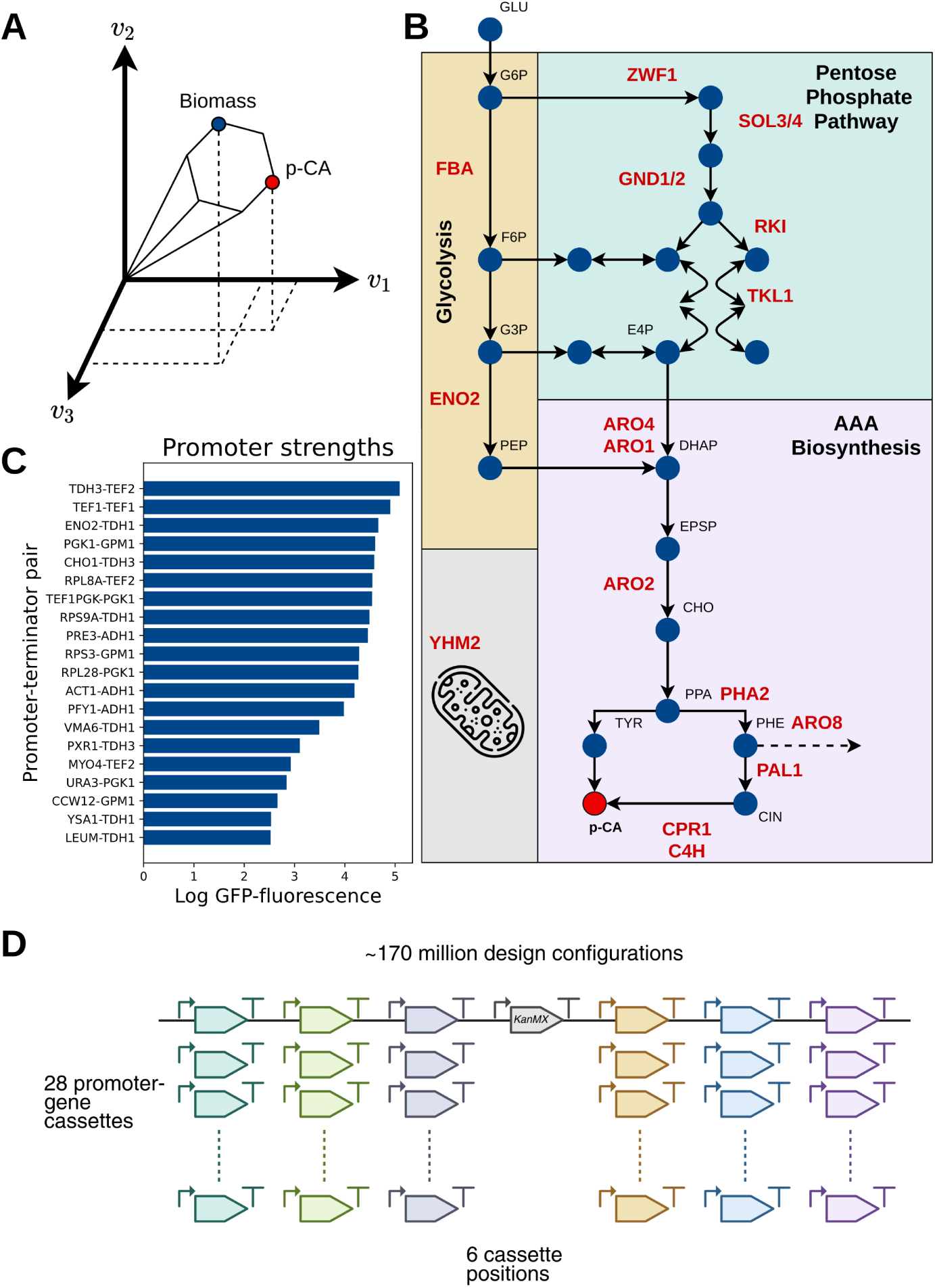
**Feature selection using the constraint-based model Yeast8**. A) The steady state flux ratio between p-CA and biomass as an objective for parsimonious flux balance analysis is used to choose potential targets (see Methods). B) A simplified schematic for the route from glucose to p- CA. In red, the gene targets that were found by pFBA are denoted. C) A set of previously quantified promoters for the library [12]. D) The designed library consists of 28 promoter-gene-terminator cassettes per library position, and 6 engineered positions. A KanMX selection marker was added (shown in grey). This results in a combinatorial action space of approximately 170 million designs, while some cassettes constructions did not succeed (18 out of 168).

Variation in the input feature values is important for the predictive capabilities of machine learning models. We therefore used twenty promoters that were previously quantified using GFP fluorescence, ranging several orders of magnitude in intensity [12] (Fig. 2C). This resulted in a library design with 28 promoter-gene cassettes, totaling approximately 170 million design configurations.

### 2.2 Supervised modeling of the p-CA pathway to predict strain designs

Despite increasing capabilities of high-throughput screening and sequencing, it is both infeasible and undesirable to exhaustively sample all combinatorial strain designs. Successful strain recommendation for follow-up DBTL cycles requires a well-performing supervised model that can predict unseen designs [23]. o train this machine learning model, a library transformation was performed followed by screening of 3000 randomly picked colonies and subsequent measurement with Nuclear Magnetic Resonance (NMR). p-CA production was plate-corrected and normalized with respect to the parent strain (SHK0066) [12]. We observe a large spread in p-CA production between strains, with 5% of the strains producing more than twice compared to SHK0066 and 27% produced less than SHK0066 (Fig. 3A). To ensure that the top producers were indeed high performers, we re-screened the top 86 with four biological replicates. This resulted in a more conservative estimate of the p-CA production, but many remain to be high producers (see SI Rescreening and rebuild of top strains). These remeasured top producers, as well as data from a previous DBTL cycle were included in the training data [12].

**Fig. 3:**
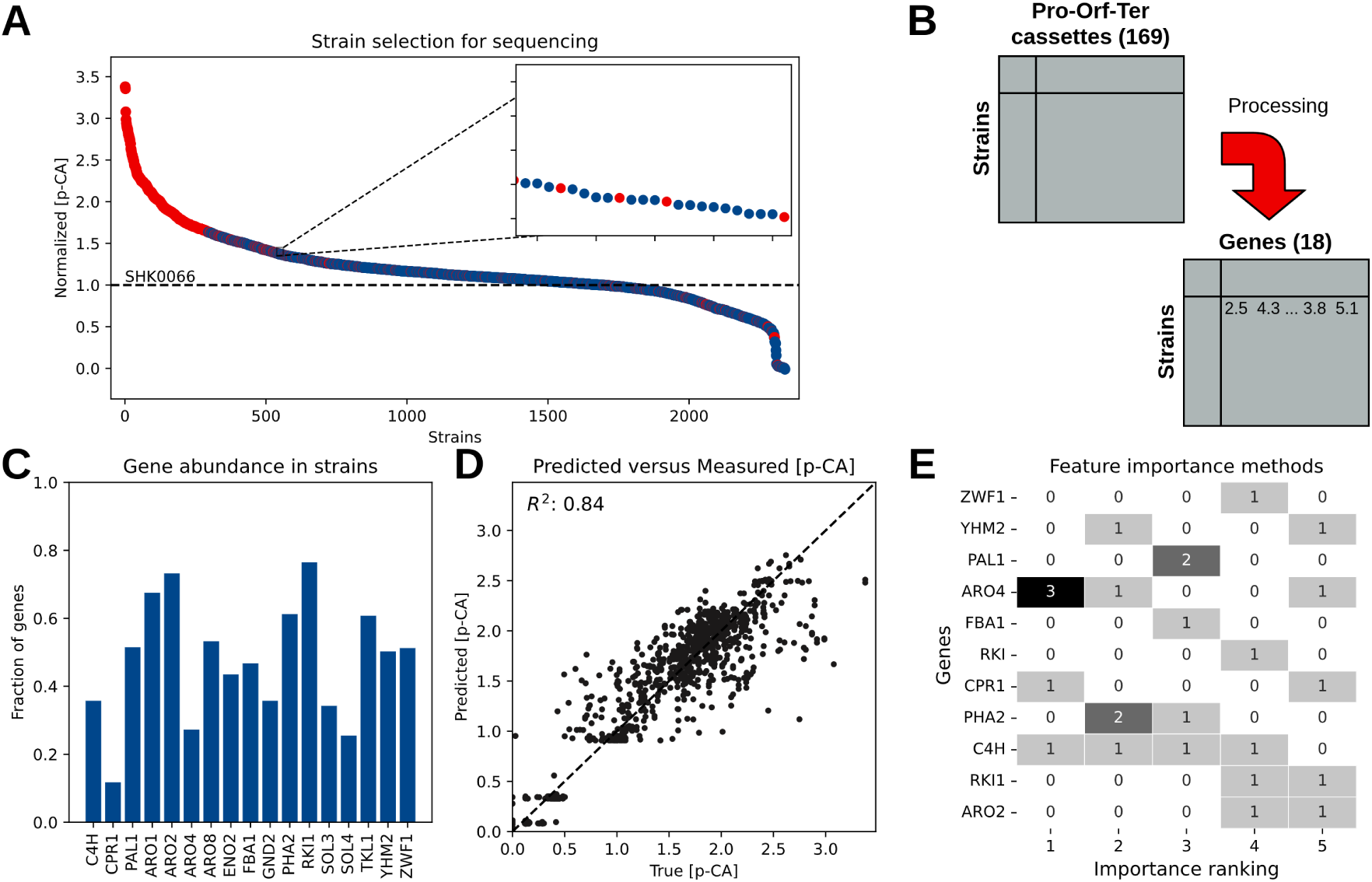
Screening, sequencing and processing for supervised machine learning. A) Strains selected for screening and sequencing. 3000 colonies were screened for p-CA production, of which 2350 strains passed screening quality checks. Of the 2350 strains, six 96-wells plates were chosen for sequencing (shown in red). These consist of the top producers, but also a representative sample of lower producing strains to mitigate biases for machine learning using XGBoost[19] B) After sequencing and several preprocessing steps (see Methods), the count matrix for the cassettes is processed to a gene numeric matrix, where values are the GFP quantified promoter strengths. C) Gene abundance in the strains that were sequenced. The fraction indicates the abundance of the gene in the total set of strains. Gene abundance in the strains that were sequenced. The fraction indicates the abundance of the gene in the total set of strains. All library genes appeared in the strains. D) Predicted versus true p-CA production for the test folds in cross-validation. A high Pearson Correlation Coefficient is observed, indicating that the processing leads to a good representation for p-CA prediction. E) A count table of the ranking of five feature importance methods (see Methods). The count is the number of times a particular gene was found to be important in that rank. While there is some consensus between the methods, much variation exists in the resulting rankings.

With a library transformation, the strain’s genetic composition is not known after screening. We therefore chose a representative set that covers the production range and performed whole genome sequencing to determine the integrated library cassettes (see Methods) (Fig. 3A). Further processing of the resulting copy count matrix was performed by collapsing cassettes with the same genes to numerical values determined by the promoter strengths, which led to effectively reducing the dimensionality of the dataset from 169 library cassettes to 18 genes (Fig. 3B). Overall, we observed that all genes appeared in the sequenced strains with high abundance (*>* 10%), with the exception of CPR1 (Fig. 3C). This is however in line with the number of positions that CPR1 was included in during library design, with only five out of the 150 succeeded cassettes having CPR1 as an open reading frame. Although the library was designed to introduce six genes in each of the strains, undesired homologous recombination resulted in strains with up to 20 introduced genes (see Dis- cussion). These strains might be genetically unstable but still contain valuable information for the machine learning model and were thus included in the training dataset.

For modeling, the ensemble method XGBoost with a density based weighted loss function was used, which weights underrepresented data-points higher using kernel density estimation [24] (see Methods). We validated model predictions by cross-fold validation. The predicted versus true p-CA production shows a high Pearson correlation on the validation sets (PCC = 0.84), which suggests a strong overall predictive performance. However, some under-prediction for high-performing strains, as well as some over-prediction of poorly performing strains is observed (Fig. 3D). To further validate the model, we aimed to understand which genes were contributing to model performance and high p-CA production. Five feature importance methods were used to identify gene importance rankings (see Methods), which resulted in eleven out of the eighteen genes to be found by at least one method (Fig. 3E). Three genes (ARO4, PHA2, C4H) were found by more than three methods and were also reported to increase p-CA production [12, 25, 26]. Overall, the gene importance rankings found based on this model are in line with previous research, despite the variability between the results of each individual method. The strong predictive performance of the model in combination with the gene importance rankings that align with known biology suggests the model’s reliability for further use.

### 2.3 A model-based exploration-exploitation strategy for strain recommendation

Along with the trained and validated XGBoost model, a strain recommendation strategy to improve p-CA production is required to navigate the vast combinatorial design landscape. This recommendation strategy needs to define an action space that is predicted by the model, but is also feasible from a metabolic engineering perspective (Fig. 1B). Furthermore, it requires a method to deal with uncertainty of model prediction. Despite the high performance on the cross-validated sets (Fig. 3D), it is typically observed that predictive performance drops when testing new strain designs. One reason for this is large biological/technical variation in screened p-CA concentration, especially for high-performance strains (see SI Rescreening and rebuild of top strains). Another reason may be model overfitting. A more fundamental reason for this drop in performance, however, is that often a new parent strain is used for a follow-up DBTL cycle. Recommendations made for the new parent are therefore outside of the training data domain, a domain where machine learning models typically struggle. The balance between exploiting the model predictions and exploring the space of designs for potentially higher rewards is therefore important [10]. For these reasons, a recommendation strategy that diversifies the set of recommended strains by balancing exploration and exploitation are crucial for a successful pathway optimization strategy. Here, we introduce such a model-based recommendation strategy with a balancing of exploration/exploitation.

We construct a design space by pairing four promoters with eighteen genes, resulting in 72 cassettes (Fig. 4A). For each pathway design, four gene cassettes were chosen (see Methods). The p-CA production of these *in silico* strain designs is predicted by the model described above. The softmax distribution is then used as a method to sample these actions, which assigns a sampling probability to each action given the predicted p-CA production. The exploration-exploitation parameter *β* shown in the formula of figure 4A controls the magnitude of the bias in the sampling distribution: the higher *β*, the stronger the sampling will be biased to exploit the model predictions for recommending high-producing strains. To highlight the consequences of increasing the exploration-exploitation parameter, 25 strains were sampled from this softmax distribution with increasing *β*. The number of unique genes in the sampled design set shows a decreasing trend with increasing exploitation (Fig. 4B), while the average and maximum strain performance increases (Fig. 4C). This shows that the set of sampled strains becomes more similar in highly exploitative scenarios, which may be problematic when the underlying supervised model is biased. There is therefore a clear trade-off between the gene diversity in the set of sampled strains and the average predicted production of this set (4D).

**Fig. 4:**
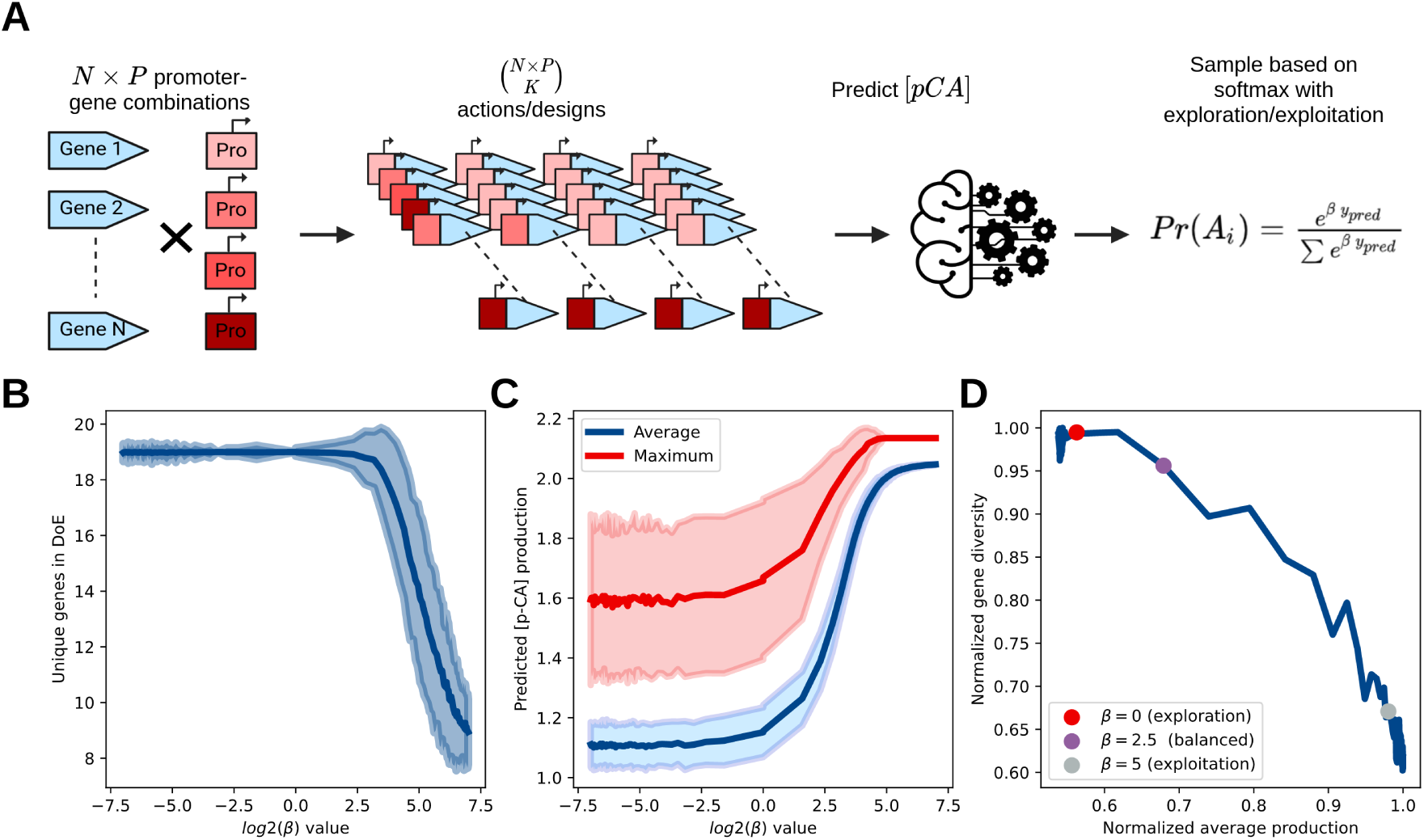
Machine learning-assisted automated recommendation with exploration/exploitation balancing. A) Overview of the recommendation strategy. A set of four promoters (*P*) with eighteen genes (*N*) is constructed and based on the number of positions in the pathway (*K*), the action space is constructed. Then, the model reported above is used to predict p-CA outcomes. The designs that will be built are sampled from the softmax distribution that has an exploration-exploitation parameter (*β*). B) Influence of the exploration/exploitation trade-off on diversity in a set of designs that were sampled. Twenty-five samples were taken per *β* value, and this was repeated 100 times (shaded region). As *β* increases, the gene set diversity goes down. C) Average and maximum predicted production as a function of *β*. As exploitation increases, the average predicted production increases. D) The trade-off between gene diversity in the samples and average production. Three *β*-settings were chosen for experimental validation of the recommendation algorithm and to experimentally observe the influence of this parameter.

### 2.4 Experimental validation reveals a successful automated recommendation strategy and importance of balancing exploration-exploitation

The exploration-exploitation trade-off is a central concept in Bayesian optimization and reinforcement learning applications [10, 27]. Despite that this trade-off is also observed in the recommendation strategy described in the previous section and has been noted to be of importance in the context of metabolic engineering [11, 14, 15], this remains to be experimentally verified. To verify whether the exploration-exploitation trade-off is of importance for a successful recommendation strategy, we tested four scenarios with differing exploitative-ness. Three scenarios are considered for SHK0066: an explorative, balanced and exploitative scenario (respectively *β* = [0, 2.5, 5]) (Fig. 4D). While in iterative optimization the parent strain often changes between DBTL cycles, we also chose a fourth scenario, where the best predicted strain with four extra integrated genes (SHK0073) was chosen as a parent strain with a balanced exploration-exploitation strategy.

Twenty-five strains with four biological replicates were built per scenario (fig 5A). The dashed lines show the parent SHK0066 and the best performing strain when using a greedy recommendation strategy for six included genes (see 4). The mean production of the strains with parent SHK0066 increases with increasing exploitative-ness. Several strains in the balanced and exploitative scenario outperform the greedy recommendation strategy, suggesting that using predictive models in a purely fashion must be done with caution. The balanced scenario shows the highest variance of the three scenarios, while the exploitation scenario shows the lowest variance in terms of p-CA production. This high variance in the balanced scenario is expected: the set of strains with a higher genetic diversity result in a larger range of p-CA production than strains with a similar genetic makeup. These results experimentally verify that there is a clear trade-off in this recommendation strategy.

For the final scenario, where the top producer SHK0073 with four included genes was used, the mean p-CA production is even higher, with several strains strongly outperforming the greedy recommendation strategy. The best strain produces 137% more than SHK0066, 33% more than the greedy strategy (six genes), and 18% better than the best strain in the initial library transformation round (ten genes integrated) (see fig. E3B). Note that there is an even higher variance in production than in the balanced scenario of SHK0066. The higher variation in production performance for this scenario could be explained two-fold: 1) high variation is generally expected in this balanced scenario, and 2) the model was not specifically trained on this host strain. The latter argument stresses the importance of including exploration in an approach: there is no guarantee that a model that is trained on a previous strain generalizes well out-of-distribution, a fundamental limitation of many machine learning approaches. It is thus crucial to address this type of uncertainty when using automated recommendation strategies.

### 2.5 Deeper validation of predictive-ness and feature importance

Although the described recommendation strategy shows to be succesful, some questions remain on predictive performance and explainability of the model. The predictive performance is of importance in that a better predictive model will allow for a more exploitative recommendation strategy. Explainability is important from the perspective of gaining insight into the biological process, but also to gain a deeper trust that the model has learned a representation that is based on biology [28].

When we further validate predictive performance in terms of the predicted versus true p-CA production, we observe that the general trend of the scenarios is predicted well (Fig 4B). There is a high correlation observed between the predicted and true p-CA across scenarios, with a slight tendency to overestimate the p-CA concentration. However, it is also observed that predictive performance is relatively poor within scenarios (Table 1). For the exploratory and balanced scenarios (top strain SHK0073 and original parent SHK0066), we observe that variation is captured better than for the exploitative scenario. This is likely due to the low variance of the exploitative scenario, which leads to lower correlations when the prediction is off. These results indicate that the machine learning model captures the overall behavior well, but that exploration that results in higher variation of the designs is essential, as these models may overfit/overestimate.

**Table 1:** Correlations of the predicted versus true p-CA production of. **figure 4B**.

| Scenario | Pearson Correlation | Spearman Rank Correlation |
| --- | --- | --- |
| All | 0.63 | 0.78 |
| $\beta = 0$ (SHK0066) | 0.28 | 0.43 |
| $\beta = 2.5$ (SHK0066) | 0.15 | 0.32 |
| $\beta = 5$ (SHK0066) | 0.02 | 0.16 |
| $\beta = 2.5$ (SHK0073) | 0.55 | 0.76 |

To validate whether the model’s feature importance results can be experimentally verified, we performed two experiments. The first experiment consists of the integration of individual promoter-gene cassettes in SHK0066 to validate the results found by the feature importance rankings in figure 3E. PHA2, ARO4, PAL1, and C4H had a positive effect on p-CA production (t-test, p-val<0.05) (Fig. 5C). These genes were all found by at least two of the feature importance methods. While the rankings per method may not be consistent, this still suggests that the representation learned by the model is aligned with the underlying biology. The second experiment consists of a greedy recommendation, where the best predicted strain was built with an increasing number of genes (Fig. 5D). While a steady increase in production is observed with the increasing number of included genes, the improvement was overestimated by the model. The best strain using this greedy approach was shown to produce lower than many strains proposed by our recommendation algorithm. This suggests that one has to be careful with using a purely greedy approach when building strains.

**Fig. 5:**
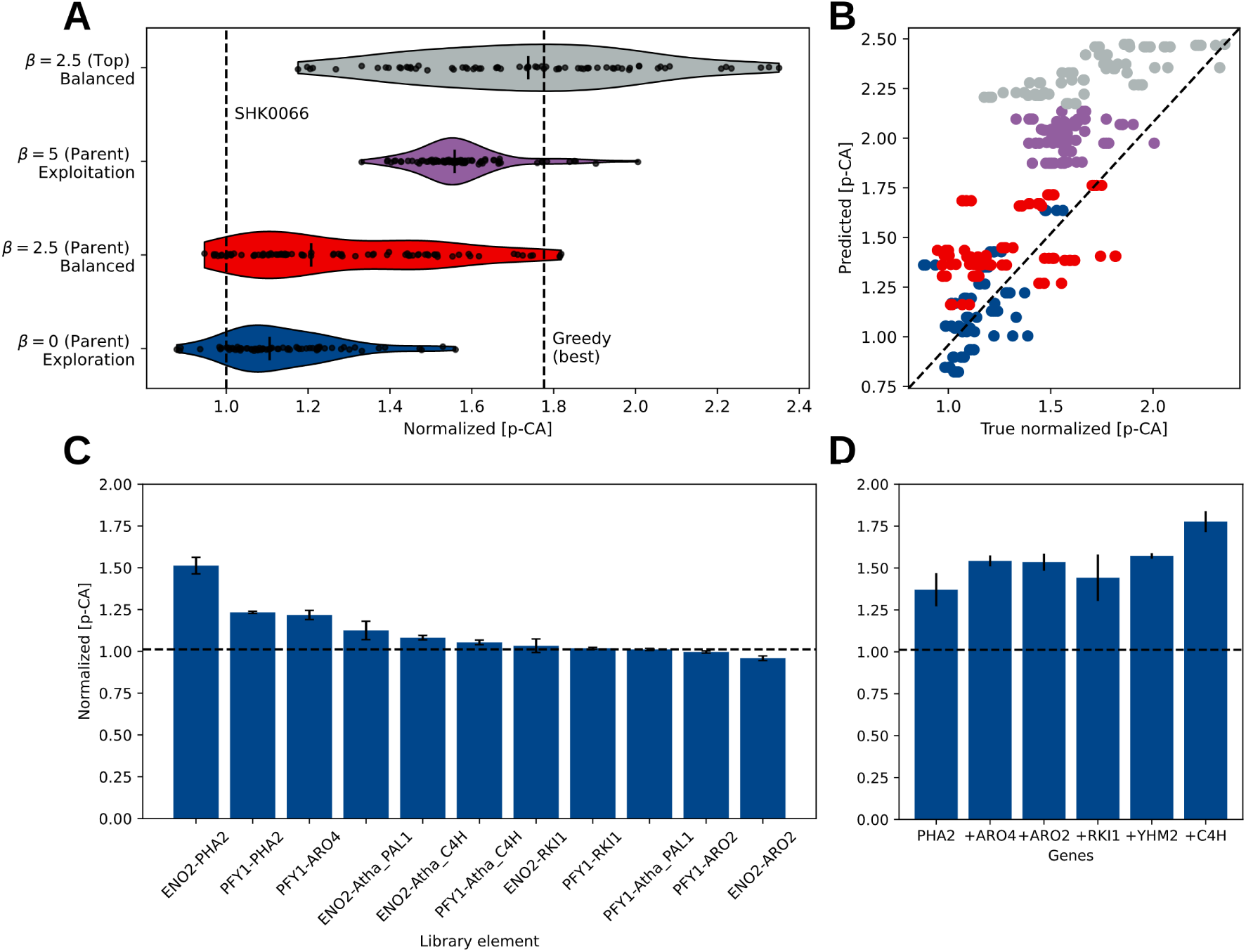
E**x**perimental **validation of the effect of balancing exploration-exploitation in large combinatorial spaces.** A) An explorative, balanced, and exploitative scenario (*β* = [0, 2.5, 5]) was tested for parent SHK0066. For the top predicted strain (SHK0073), a balanced scenario was tested. B) Predicted versus true [p-CA] production for the four scenarios. XGboost predicted the overall trend well, but variance in the actual [p-CA] production was underestimated. C) The experimental effect of individual genes that were included in the library design. A weak (PFY1) and strong promoter (ENO2) were used. PHA2, ARO4, C4H, and PAL1 have a significant effect on p-CA production (Wilcoxon rank sum test, *p <* 0.05). D) Greedy strain building based on the best predicted strain by the model. Six strains were built with an increasing number of cassettes that were predicted to maximally increase production.

## 3 Discussion

This work explores the potential of automated recommendation tools for metabolic engineering given a large combinatorial space that cannot be explored experimentally. By expanding the combinatorial space up to approximately 500 million theoretically possible designs, the need for exploration instead of exploitation becomes more prominent. We show that successful recommendation using a balanced exploration-exploitation of the combinatorial space outperforms greedy approaches that are purely based on model predictions. We also observe that a high-performing parent strain combined with a balanced recommendation scenario outperforms in terms of p-CA production. This is to an extent surprising, as there is no guarantee that a machine learning model that predicts action outcomes is able to predict outside of the training data distribution [29].

The combinatorial design space considered in this study was constructed from eighteen genes and twenty promoters, with six library positions and 28 cassettes per position. This means that there is a large repetitiveness in the library design, which consequently leads to many strains with homologous recombination events. Some strains were shown to have over ten included genes instead of the designed six, which makes them genetically unstable. For example, the top two strains in the first DBTL cycle had 26 and 23 genes (including the KanMX markers) integrated, respectively (Fig. Rescreening and rebuild of top strainsB). Despite the fact that these strains may not be genetically stable, they were still informative for training the model and were therefore included during training of XGboost.

An important aspect of this work is the importance of balancing exploration/exploitation of the combinatorial design space. As far as we are aware, this is the first study that experimentally verifies this effect in the context of metabolic engineering, although several studies have acknowledged its importance [13, 15, 30]. While the exploitative scenario (*β* = 5) showed high performing strains, this comes at the expense of genetic diversity of proposed designs. This may result in choosing suboptimal actions when model performance is less compared to the p-CA model used in this study. It is therefore essential to take model uncertainty into account when balancing exploration/exploitation of the design space. While we show that this balancing is important, we did not propose a direct approach to how this should be done. One way to balance exploration/exploitation may be to maximize the product of the genetic diversity and average production of figure 4D, similar to the approach taken in the MODIFY algorithm for directed evolution [31].

Here, we chose for the second round to build specific strains that were recommended by the algorithm. As the methodology to build these strains differs from a library transformation, the number of strains that can be build is significantly smaller than what can be achieved with a library transformation given comparative experimental resources. However, this recommendation strategy could similarly be applied to generate library designs by using the approach shown in figure 4A. An additional step is then required to generate the probability distribution per library element per position. This can then be experimentally implemented by changing the concentration of library cassettes, which change the probability of being integrated during library transformation. We have experimentally verified that concentration changes can be used to modulate the chance of cassette integration in the supporting information (Supporting information Whole genome sequencing: library pools). This would allow for larger follow-up rounds.

To test feature importance methods, we experimentally determined the individual influence of genes on p-CA production (Fig. 5C). The four genes (PHA2, ARO4, PAL1, C4H) that were important were also found by the feature importance methods, with mixed results on their relative importance. PHA2 catalyzes the conversion of prephenate to phenylpyruvate [32], and was reported to have a significant effect on p-CA production [33]. ARO4 is the first reaction in the shikimate pathway and was already considered to be important for the used parent strain in this study [12], as well as in another p-CA yeast cell factory [34]. PAL1 and C4H were genes that originates from *Arabidopsis thaliana* and provides an alternative route to the TAL-route [33, 34]. These genes were also previously included in the parent strain [12]. These four genes were all found to be important by the five feature importance measure we used, but some other targets were also found in the importance ranking (Fig. 3F).

## 4 Methods

### 4.1 Organisms and media

S. cerevisiae strains originate from CEN.PK113-7D. Coumaric production strain BMP+PAL-C4H- CPR (SHK0066) from study [12] was used as starter strain. Strains were grown at 30°C in Yeast Extract Phytone Dextrose media (YEPhD, 2% Difco™ phytone peptone (Becton-Dickinson (BD), Franklin Lakes, NJ, USA), 1% Bacto™ Yeast extract (BD) and 2% D-glucose (Sigma Aldrich, St Louis, MO, USA)). When required, antibiotics were added to the media at appropriate concentrations: 200 µg/ml nourseothricin (Jena Bioscience, Germany), 200 µg/ml geneticin (G418, Sigma-Aldrich). E. coli DH10B (New England BioLabs, Ipswich, MA, USA) was used as cloning strain and grown at 37°C in 2*Peptone Yeast Extract media (2*PY, 1.6% tryptone peptone (BD), 1% Bacto™ yeast extract (BD) and 0.5% NaCl (Sigma Aldrich)). When required, antibiotics were added to the media at appropriate concentrations: 100 µg/ml ampicillin (Sigma-Aldrich), 50 µg/ml neomycin (Sigma-Aldrich). Solid medium was prepared by addition of Difco™ granulated agar (BD) to the medium to a final concentration of 1.5% (w/v).

### 4.2 Cassette construction

#### 4.2.1 Gene Targets

To determine which gene targets should be included in the library, parsimonious flux balance analysis (pFBA) is performed [22] on the genome-scale model Yeast8 [21]. We use the pFBA solution of a p-Coumaric acid objective and the biomass objective to calculate a flux ratio. This flux ratio indicates which fluxes are rerouted when diverting carbon flux from biomass towards p-Coumaric acid and was used to select genes for further validation (*v^ss^*). After excluding sink, exchange and non-targetable reactions, 33 potentially interesting gene targets were left.

We further aimed to substantiate gene targets by performing literature enrichment analysis using the STRING database [35], thermodynamic information from the equilibrator database [36], and inhibitor information from the BRENDA database [37]. Finally, we assess the importance of each gene by literature review (see table SI1). This resulted in a set of nineteen validated genes that are used in the library. Open reading frames (ORF) were codon pair optimized for *S. Cerevisiae* [38]. Cassettes were designed as described in [39] and constructed as described in [12]

#### 4.2.2 Promoters

A set of twenty previously established promoters with varying strengths quantified through GFP-fluorescence were used in the library design [12]. The expression strength of these promoters spans several orders of magnitude and can be used to regulate gene expression. We assume that the expression pattern might differ between genes for the same promoter, but that the relative expression per gene is similar.

#### 4.2.3 Library design

The number of positions in the library was limited to six factors to ensure sufficient transformation efficiency. To not exclude certain gene combinations during transformation, we decided that for each position in the construct all genes should be present with two distinct promoters [12]. For genes that were previously considered in the reference strain (C4H, CPR1, PAL1, ARO4), we chose one promoter per library position. This library consists 168 cassettes, of which 150 were successfully build. While incorporating all genes ensures that we do not exclude certain combinations, it might lead to increased homologous recombination events during transformation. The library design can be found in the supporting information.

### 4.3 S. cerevisiae strain construction

#### 4.3.1 Transformation

Strains were constructed as described in [12]. In short, host strain SHK0066, pre-expressing Cas9 was used for CRISPR mediated integration of the cassette library into the target locus INT28, see supplementary info for gRNA design and genome locus position. Cassette libraries were prepared with equimolar amounts of cassettes (100ng/kb). Per cassette library previously considered genes (C4H, CPR1, PAL1, ARO4) were added with higher dosage as follows; library 1: 300ng/kb, library 2: 200ng/kb, library 3: 100ng/kb, library 4: 50ng/kb, the library design can be found in the supporting information. Linear gRNA (210ng/kb) and linear backbone with homologous sites to the gRNA (35ng/kb) were transformed together with the cassette libraries into the host strain following LiAC/ssDNA/PEG method [40].

For design-based transformation, the cassettes were prepared per pathway design in equimolar amounts and transformed with the linear gRNA and linear backbone following the same LiAC/ss- DNA/PEG method. The cassette design with on five prime and three prime side 50 base- pair connector sequences homologous to INT28 facilitate in vivo recombination of cassettes into the genome.

The transformed cells were plated on a Qtray (Nunc) containing yephD agar and appropriate antibiotics. 48-dividers (Genetix) were used in the Qtray when a pathway design was done. Qtrays were incubated for three days at 30°C and picked with Qpix 450 (Molecular Devices) into 96 MTP (Nunc) containing YephD agar with appropriate antibiotics.

#### 4.3 p-CA screening in *S. cerevisiae*

##### 4.4.1 Strain fermentation

p-Coumaric acid production experiments were done as follows; colonies were grown in microtiter plate (MTP) containing YephD agar medium and grown for 48hrs at 30°C, 750 rpm, 80% humidity. Cultures were inoculated in half deepwell plate (HDWP, thermofisher scientific, AB-1127) containing 350*µ*l YephD and appropriate antibiotics as a preculture. Strains were reinoculated in main culture into HDWP containing 350ul Verduyn Luttik with 2% glucose [41] and incubated for 48hrs at 30°C, 750 rpm, 80% humidity. Every plate contains a medium control and 16 times the parent strain SHK0066 for later normalization.

##### 4.4.2 p-CA measurment with NMR

For Flow-NMR measurements 250*µ*l end of fermentation broth was sampled into a 96-deepwell plate and mixed with 500*µ*l acetonitrile using the EPmotion (Eppendorf) to start cell lysis. The samples were centrifuged at 4000rpm for 10 minutes and 500 *µ*l supernatant was sampled into a new deep well plate. 70% of the liquid was evaporated and 100*µ*l of 1 g/L internal standard 1,1- difluoro-1-trimethylsilanyl methylphosphoric acid (FSP, Bridge Organics) was added. The samples were lyophilized to remove non-deuterated solvents. To the lyophilized samples, 600 *µ*l of D2O (Cambridge Isotope Laboratories (DLM-4)) was added and homogenized. The samples were analyzed on a CTC PAL3 Dual-Head Robot RTC/RSI 160 cm robotic autosampler (CTC Analytics AG, Zwingen, Switzerland) fluidically coupled to a Bruker spectrometer Avance III HD 500 MHz UltraShield [26]. 1H spectra were recorded with standard pulse program (zgcppr) with following parameters: 16 scans, 2 dummy scans, 33k data points, 16.4 ppm spectral width, 1.2 s relaxation delay (d1), 8 *µ*s 90° pulse, 2s acquisition time, 15 Hz water suppression, and fixed receiver gain (rg) of 64 Spectra were processed and analyzed using Topspin 4.1.4 (Bruker). Spectral phasing was applied and spectra were aligned to 3-(trimethylsilyl)-1-propanesulfonic acid-d6 sodium salt (DSS- d6, Sigma-Aldrich) at 0 ppm. Auto baseline correction was applied on the full spectrum width. Additional third-order polynomial baseline correction for selected regions was applied if needed. The amount of pCA (doublet, 6.38 ppm, n=2H) was calculated relative to the signal of FSP [12].

#### 4.5 Strain quality control using whole genome sequencing

##### 4.5.1 Choice of screened strains for sequencing

Based on the screening of 3000 strains, we chose a representative set of strains for sequencing. We chose three 96-wells plates for the top producers, two 96-wells plates for the strains that showed improved p-Coumaric acid production but that are not in the top, and one 96-wells plate for strains that performed worse than the parent strain.

##### 4.5.2 Whole genome sequencing

*S. cerevisiae* cells (OD600 of 5-10) were pelleted and lysed as follows; 140mg Zirconia beads (0.5mm diameter, Biospec products) and 400*µ*l ZymoBIOMICS Lysis buffer (Zymo Research) were added to the cell pellet. Cells were lysed for 5 minutes at 1500 rpm with the Geno/Grinder 2010 (Cole-Parmer).

gDNA was further isolated following ZymoBIOMICS 96 DNA kit (Zymo research) according to manufacturers protocol. The isolated genomic DNA was quantified using the Tapestation 4200 (Agi- lent) with a genomic DNA screentape (Agilent) and Nanodrop (Thermofisher Scientific). Genomic DNA was sequenced per strain or per library pool of transformants using the native barcoding kit (SQK-NBD114.96) from Oxford Nanopore Technologies on a Promethion device (FLO-PRO114M) according to manufacturer’s protocol. For pool sequencing super-accuracy basecalling was used and for strain sequencing high accuracy basecalling.

##### 4.5.3 Data processing of individual clones

DNA sequencing reads obtained from picked individual colonies after transformation, were cleaned using Porechop [42]. Cleaned reads were then assembled into genomes using Canu [43], with the genome size parameter set to 12 Mbp, and were followed by one round of polishing by Racon [44] and Medaka [45]. To identify where the inserted BioBricks were located, annotations of the cassette sequences (such as promoter, open-reading-frame, terminator annotations) were transferred to the genome assemblies by blast-based sequence alignment, leading to genome assembly annotations in GFF format. Cassette annotations were only transferred to the genome assemblies if at least 95% of the cassette annotation was included in an alignment hit to the genome, with at least 95% identity.

#### 4.6 Machine learning-assisted recommendation strategy

Ensemble learning tree methods have shown to perform well on combinatorial pathway optimization data [13, 17]. We therefore trained an XGBoost [19], a regularized gradient boosting algorithm, with a modified loss function that is more specific to our purpose.

##### 4.6.1 Density-based weighting of loss function

Machine learning methods typically assume that data is balanced; that is, the target variable is uniformly distributed. However, in production screening data this is often not the case. High performing strains are typically underrepresented, while strains with minor increases in performance are overrepresented. To mitigate the effect of this imbalanced distribution on the mean squared error (MSE) [46], we used DenseWeight [24]. DenseWeight weighs the influence of data instances on the loss function according to the target variable frequencies estimated through kernel density estimation. DenseWeight requires setting two hyperparameters that determine how much the density estimation should influence the weighing and was set to *α* = 0.5 and *ɛ* = 0.01.

XGboost requires calculation of the gradient and hessian of the loss function for efficient training and due to the modified loss function needs to be modified. The modified loss function (*L_DensLoss_*) is the mean squared error weighted by a function (*f_w_*(*y_i_*)) that is dependent on the target variable density and its hyperparameters. The loss function, Jacobian, and Hessian with respect to a data instance *y_i_* is then (eq. 1-3).

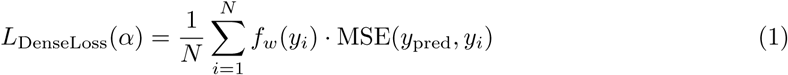

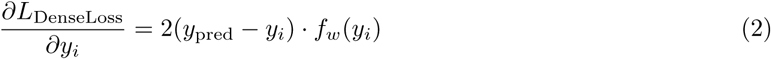

##### 4.6.2 XGboost training

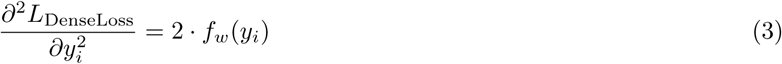

Bayesian hyperparameter optimization was performed on the training data using scikit-optimize [47]. This resulted in the following hyperparameters (*reg_lambda=1*, *max_depth=2*, *tree_method=”auto”*). The customized loss function was used as described above. 200 rounds of boosting were performed with an early stopping criterion of 40. This means that if the validation mean squared error does not improve with 40 rounds, training is stopped. Model validation was performed on a ten percent random split using sklearn and evaluated with the Pearson correlation coefficient between true and predicted. We further validated the model using an external validation set of designs (see below).

##### 4.6.3 Recommending new strains using gradient bandit principles

To recommend new strains for the second round, we start by defining the action space (Figure 1B). In the case of *N* = 20 genes with *P* = 4 cassette positions, we choose *X* = 4 different levels of promoter strengths, which results in *^N×X^* = 1028790 actions. Actions that have multiple integrations of the same gene are filtered out. Then, XGBoost is used to predict the potential outcome of the set of actions given the reference strain. We chose two reference strains: the parent of the first library transformation round (SHK0066) and the best predicted action strain for four pathway positions.

As a recommendation strategy based on predicted outcomes, we used a gradient bandit approach [10] using the softmax distribution with an exploration/exploitation (*β*) parameter (eq. 4).

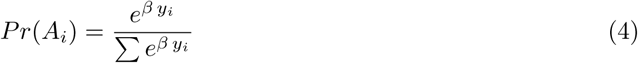

The exploration/exploitation parameter *β* is an important parameter for automated recommendation and represents a trade-off between the diversity of the set of chosen designs in terms of gene content and the expected performance of the strains in terms of production ([31]). Here, we have tested three exploration/exploitation values (*β* = 0, *β* = 2.5, *β* = 5) to explore this trade-off in more detail.

##### 4.6.4 Feature importance methods

Five feature importance methods were used for retrieving gene importance rankings from the trained model. Two methods are intrinsic to the XGBoost model, where the *weight* method ranks based on the number of times a feature is used to split data across the trees and the *gain* method bases this one the average information gain across all splits the feature is used in. The three other methods are model independent. One popular method in explaining machine learning models are the SHAP values [48]. We also tested feature importance ranking that were based on the area under the curve of gene-promoter threshold plots as used in [17]. Finally, we also performed a ranking based on a search for the best strain when predicting for *^N×P^* actions as suggested in the recommendation algorithm described above, for *K* = [1*, …,* 6].

#### 4.7 Code and data availability

All scripts used in this study can be found in (GitHub).

Acknowledgements

## Supporting information

SI_library_design_DBTL1

SI_DoE_design_DBTL2

## Appendix A p-CA strain lineage

**Fig. A1:**
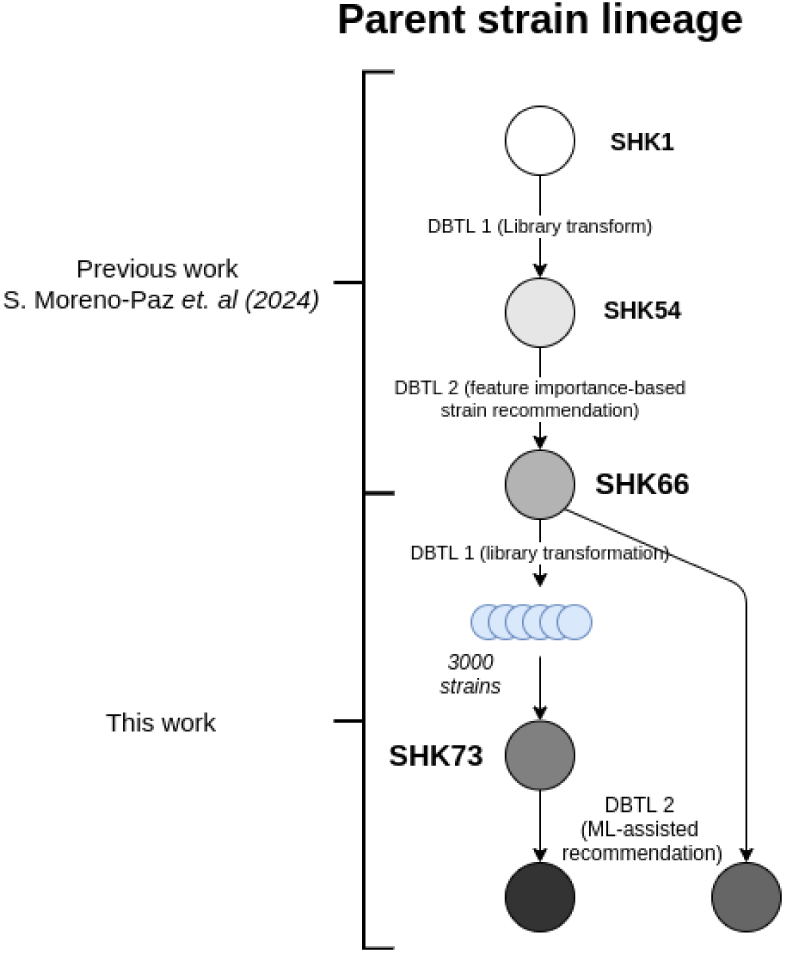
An overview of the lineage of p-Coumaric acid parent strains.

## Appendix B Library Design

### B.1 Gene target literature review

After performing pFBA with a filtering for ratio of steady-state *pCA/Biomass* flux, we gathered thermodynamic data from BRENDA, and performed a literature enrichment analysis using STRING [35, 37]. This helped us to more principally choose gene targets. On top of that, we performed a systematic literature review for the potential targets. The results are summarized in the table (Table B.1).

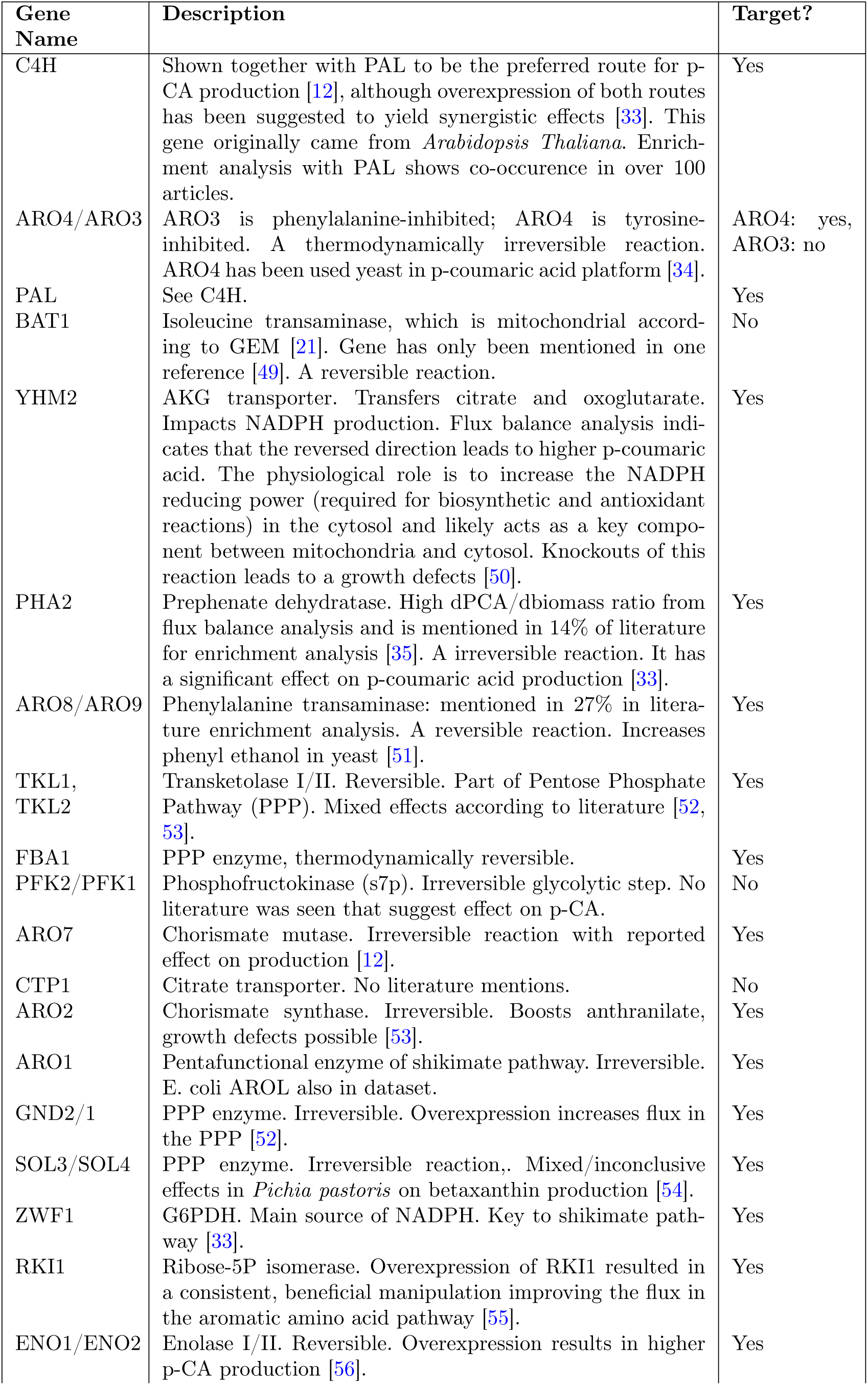
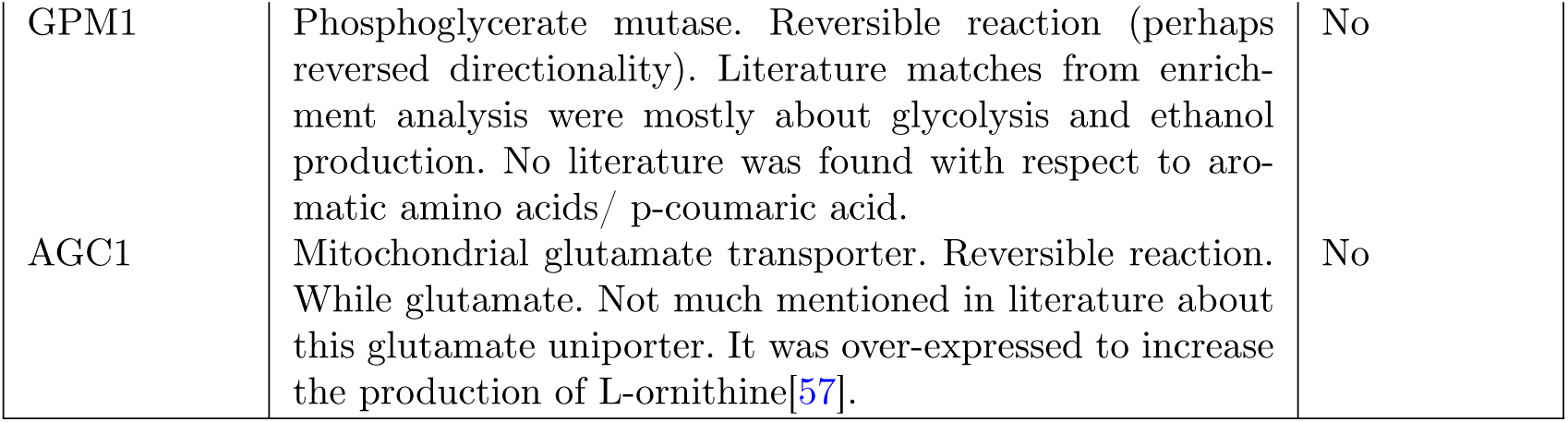

## Appendix C Library designs

Library design of the first DBTL cycle for library transformation can be found in *Round1_LibTrans_design.xlsx*. Recommended designs for the second DBTL cycle can be found in *Round2_ML_assisted_design_Strainstudio_AI4bioDoE.xlsx*.

## Appendix D Whole genome sequencing: library pools

The current implementation of the ML-assisted recommendation strategy chooses specific strain designs for the next DBTL cycle. This limits the number of strains that can be built compared to a library transformation approach. An alternative approach to predicting specific designs is to learn a probability distribution for the library elements based on the machine learning model. This bias can then be used to favor certain library elements when being transformed by experimentally modulating the dosage of cassettes according to the probability distribution. This would allow for building many more strains in a follow-up round, which could be useful when still in an exploratory phase of metabolic flux optimization. To investigate whether this cassette dosage modulation can be used to bias integration, four libraries were designed with varying dosages for four previously considered genes (C4H, CPR1, PAL1, ARO4): library 1 (300 ng/kb), library 2 (200ng/kb), library 3 (100 ng/kb), library 4 (50 ng/kb) (see Methods). Genomic DNA is then sequenced per library pool to observe whether this biasing led to higher integration occurrence for these specific genes.

The read coverage per cassette was normalized by the average coverage along the genome. Per library position, cassettes with identical open reading frame were summed, which results in pseudo-expression value per gene per position in the library. The occurrence values were then corrected by the number of positions that were occupied by this gene. For the four modulated genes (C4H, CPR1, PAL1, ARO4), occurrence values are averaged and the are compared to the other genes in the library that had a fixed dosage (100 ng/kb) (see Methods). Results are reported in figure D2. With decreasing dosage, a clear difference in occurrence value is observed. This means that by biasing the dosage of library elements, cassette integration probabilities can be modulated during transformation. While the occurrence described here is only a proxy for the frequency of cassette integration, it provides strong evidence that this can be used to bias DNA libraries. Future research could aim to address how ML-assisted recommendations could be used to design libraries directly.

## Appendix E Rescreening and rebuild of top strains

**Fig. D2:**
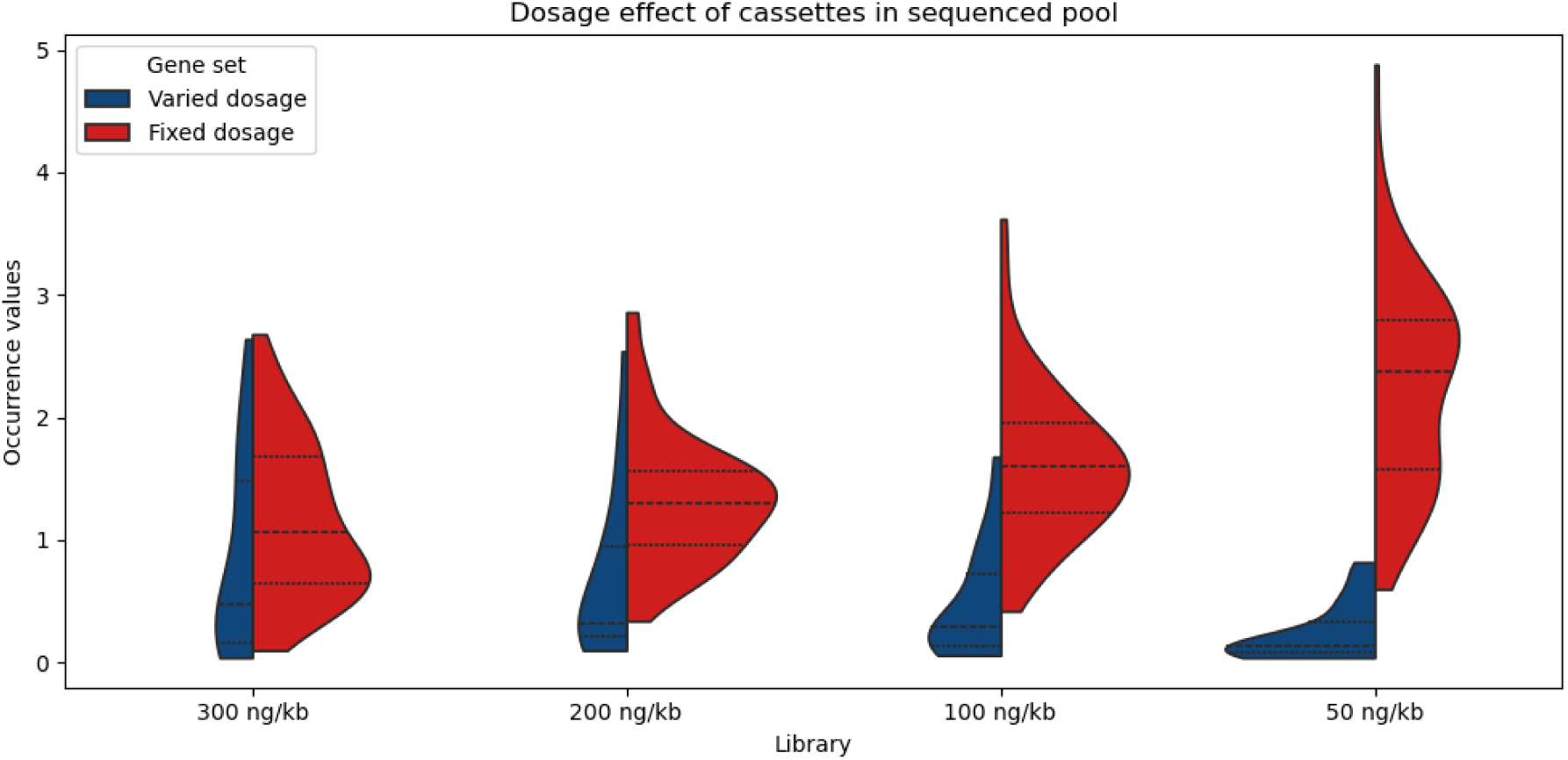
**Occurrence between the adapted dosage genes and fixed dosage genes for the four libraries**. The occurrence values are a proxy of the frequency of being integrated into strains during library transformation. The varied gene set consists of C4H, CPR1, PAL1, ARO4, and the fixed set consists of ZWF1, YHM2, TKL1, SOL3/4, RKI1, PHA2, GND1/2, FBA1, ENO2, ARO8, ARO2. A significant difference between occurrence values was observed for all four library designs (Wilcoxon rank sum test). With decreasing gene dosage, these genes are less represented in sequenced pool. This suggests that dosage can be used to bias libraries during transformation.

**Fig. E3:**
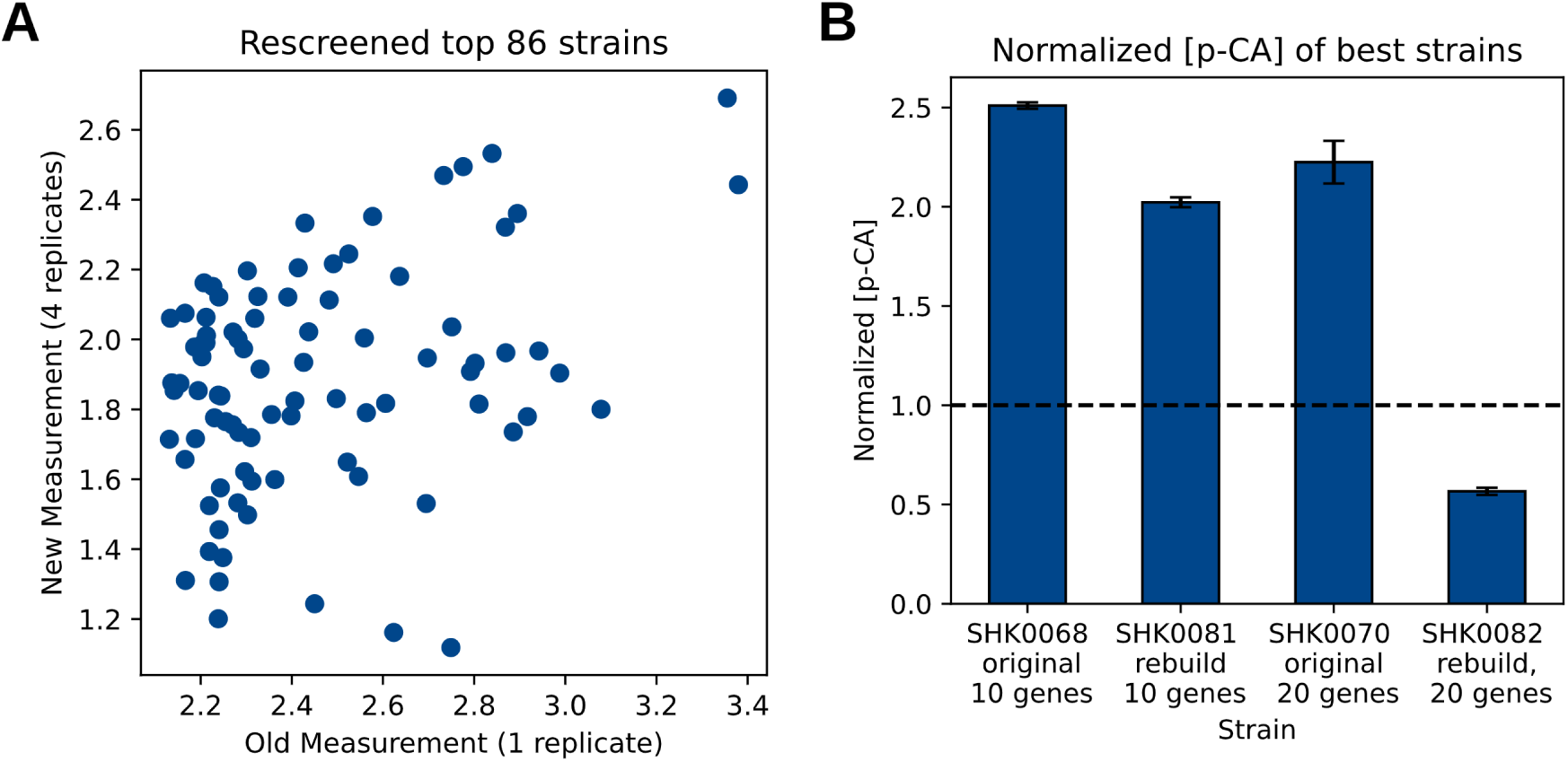
Rescreening of the top 86 strains and rebuilding of the best two strains. A) The top 86 strains were rescreened for p-CA production with four technical replicates (y-axis). The re-screened measurements show a more conservative estimate of the p-CA production. B) The two best strains in the library transformation, original measurement and the rebuilt version. Both strains had homologous recombination events, therefore exceeding the number of designed library elements in each strain.

